# A receptor-independent signaling pathway for BDNF

**DOI:** 10.1101/2022.08.23.504973

**Authors:** Julia Fath, Franck Brouillard, Alexandre Cabaye, Damien Claverie, Philippe Nuss, Victoria Poillerat, Serge Chwetzoff, Tahar Bouceba, Elodie Bouvier, Myriam Salameh, Jenny Molet, Aïda Padilla-Ferrer, Philippe Couvert, Francine Acher, Marie-Pierre Golinelli-Cohen, Gérard Chassaing, Germain Trugnan, Christophe Bernard, Jean-Jacques Benoliel, Chrystel Becker

**Author notes:** These authors contributed equally to this work (Julia Fath, Franck Brouillard, Christophe Bernard, Jean-Jacques Benoliel, Chrystel Becker). Corresponding author: Chrystel Becker, Université Paris Cité, INSERM, U1124, Équipe “Myélinisation et Pathologies du Système Nerveux”, 45 rue des Saints-Pères, 75270 Paris Cedex 06, France.

## Abstract

In addition to its well-known receptor-mediated function in cell survival, differentiation and growth, we report that the extracellular brain-derived neurotrophic factor (BDNF) also controls the intracellular KEAP1-NRF2 cytoprotective system by a receptor-independent pathway. Extracellular BDNF can cross the cell membrane as it possesses a protein-translocation domain, also known as cell-penetrating peptide. This membrane crossing process is energy-independent, ruling out endocytosis and receptor-dependent mechanisms. Once in the cytosol, BDNF binds to KEAP1 with a nanomolar affinity, enabling nuclear translocation of NRF2 and transcription of NRF2-target genes. BDNF is thus a major regulator of NRF2 activation. A dysfunction of this BDNF-KEAP1-NRF2 pathway may be involved in most diseases where antioxidant and cytoprotective functions are altered. This novel form of communication, whereby a receptor ligand protein exerts a biological activity by crossing the cell membrane, opens new avenues for cell signaling.

## Introduction

Brain-derived neurotrophic factor (BDNF), a member of the neurotrophin family, controls not only the growth, differentiation and survival of brain cells, but also synaptogenesis and synaptic plasticity ^1^. This secreted protein exerts those pleiotropic functions by binding to tyrosine kinase B (TrkB) receptors and, like other neurotrophins, to the non-selective p75^NTR^ receptor, triggering the activation of intracellular signaling pathways ^1,2^. BDNF has emerged as a molecular target for neuroprotective and functional restorative treatments in neurological and psychiatric disorders, including Alzheimer’s disease and depression ^3,4^. In stress-induced vulnerability to depression, BDNF, a biomarker identifying a population at risk of developing a depression profile, is decreased. Those low levels of BDNF are associated with a decrease in nuclear translocation of NRF2 (nuclear factor erythroid-derived 2-like 2) ^5^.

Considered as a major regulator of the cellular response to various stress factors, NRF2 contributes to the maintenance of homeostasis by regulating the expression of numerous cytoprotective genes involved in different cellular processes including inflammation, detoxification, response to xenobiotics and redox homeostasis ^6,7^. In basal conditions, NRF2 is bound *via* its high-affinity ETGE motif to a KEAP-1 (Kelch-like ECH-associated protein 1) homodimer in the cytosol ^8–10^. KEAP1 dimers form RING E3-ubiquitin ligase complexes with Cullin-3 which continuously target NRF2 for ubiquitination and proteasome degradation, thereby modulating NRF2 levels ^11^. Reactive oxygen species and electrophiles induce modifications on cysteine residues of KEAP1 leading to conformational changes that compromise its binding to NRF2 and thus promote nuclear translocation of the latter. Apart from this “canonical” mode of KEAP1-NRF2 activation by chemical inducers, alternative activation mechanisms involving cellular proteins have been described. In the so-called non-canonical pathway, intracellular proteins such as p21, p62 and dipeptidyl peptidase III (DPP3) are able to disrupt the KEAP1-NRF2 complex by direct interaction with one of the two partners, decreasing ubiquitination and degradation and increasing nuclear translocation of NRF2 ^10–12^.

Although there is an established link between BDNF and NRF2, the cascade of molecular events explaining the intracellular NRF2 activation by extracellular BDNF remains largely unknown. The latter could involve kinases downstream of its receptors ^13^ or proteins disrupting the KEAP1-NRF2 complex. In the present work, we have explored the mechanisms involved in this activation process.

## Results

### BDNF controls NRF2 nuclear translocation by a receptor-independent pathway

To investigate the receptor-dependent signaling pathway involved in NRF2 activation, we used HT-22 hippocampal neuronal cells, which express constitutively p75^NTR^ (Extended Data Fig. 1a) but not TrkB receptors (Extended Data Fig. 1b). Addition of increasing doses of extracellular BDNF, in the range of its physiological concentrations ^14^, induced a linear increase in nuclear NRF2 (Fig. 1a). This effect reached a plateau at BDNF concentrations above 7.4 nM, with nuclear NRF2 reaching ≈175% of control condition. Consistent with the increase in nuclear NRF2, cytosolic NRF2 decreased, while the levels of cytosolic KEAP1 remained unchanged (Fig. 1a,b and Extended Data Fig. 1c). Quercetin, a potent activator of NRF2 ^15^, produced similar effects (Fig. 1b and Extended Data Fig. 1c) as BDNF. Addition of a p75^NTR^-blocking antibody, at a dose preventing activation of the downstream ERK pathway ^16^ (Extended Data Fig. 1d), did not alter NRF2 nuclear translocation induced by increasing concentrations of BDNF (Fig. 1a). Quantitative analysis of transcripts shows that nuclear translocation of NRF2 is functional, since it induces the activation of NRF2-target genes involved in detoxification, drug response, catabolism and antioxidant defenses (Fig. 1c). The control of NRF2 by BDNF is bidirectional as siRNA-based downregulation of BDNF expression led to a nearly 50% decrease in nuclear NRF2 and a significant increase in cytosolic NRF2 (Fig. 1d). In NIH 3T3 fibroblasts, which do not express any known BDNF receptors ^17^, BDNF also induced nuclear translocation of NRF2 (Fig. 1e). Finally, *in vivo* BDNF microinjection into the rat hippocampus showed that the blockage of TrkB receptors and p75^NTR^ did not prevent the nuclear translocation of NRF2 induced by BDNF (Extended Data Fig. 1e). Thus, activation of NRF2 induced by physiological concentrations of extracellular BDNF is independent of any known BDNF receptors.

**Fig. 1.**
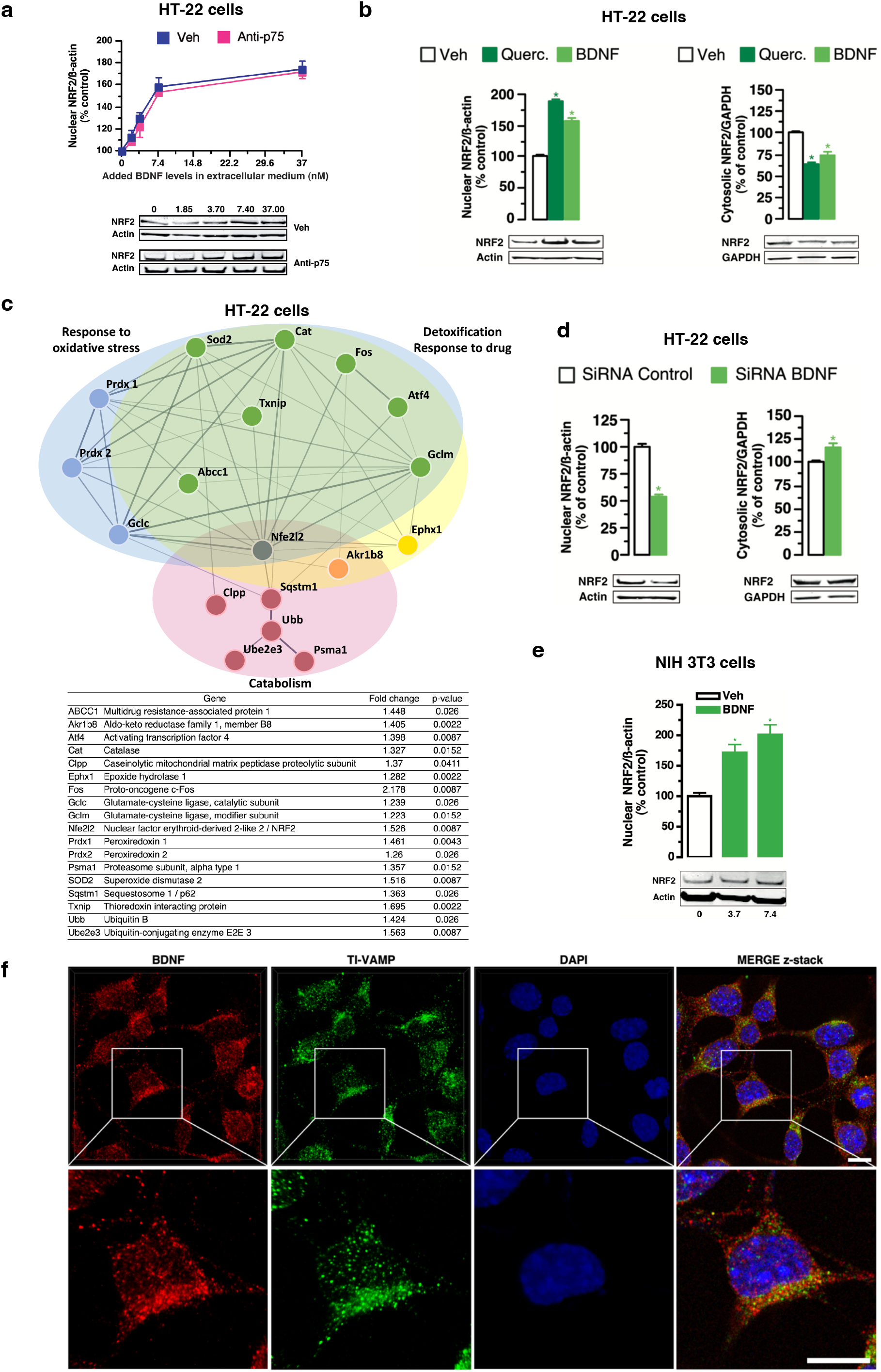
BDNF controls NRF2 nuclear translocation. **a,** Immunoblotting for NRF2 protein levels in nuclear fraction of HT-22 hippocampal cells incubated with increasing concentrations of BDNF. Incubation of HT-22 cells with BDNF leads to a linear increase of nuclear NRF2, that plateaued above 7.4 nM of BDNF (175% of control condition). The addition of a p75^NTR^-blocking antibody does not affect NRF2 translocation (*N* = 3 experiments per condition). **b,** Immunoblotting for NRF2 protein levels in nuclear and cytosolic fractions of HT-22 cells treated with quercetin (Querc., 10 μM, an activator of NRF2 as positive control), BDNF (7.4 nM) or vehicle (Veh). Incubation of HT-22 cells with BDNF or quercetin similarly increases NRF2 nuclear translocation by 60%. Application of BDNF or quercetin similarly decreases cytosolic NRF2 concentration (*N* = 4 experiments per condition). Results of statistical analyses are shown in Extended Data Fig. 5e. * *p* < 0.05 vs. vehicle (Veh). **c,** Incubation of HT-22 cells with 7.4 nM of BDNF and a p75^NTR^-blocking antibody induces the activation of NRF2 target genes, as shown by RT-qPCR experiments (*N* = 3 experiments). Activated NRF2 target genes are involved in the response to oxidative stress (blue and green), detoxification/response to drug (green and yellow) and catabolism (red and orange). **d,** Immunoblotting for NRF2 protein levels in cytosolic and nuclear fractions of HT-22 cells treated with siRNA control or siRNA BDNF. BDNF silencing in HT-22 cells results in a decrease in NRF2 nuclear translocation and an increase in NRF2 cytosolic levels (*N* = 3 experiments per condition). * *p* < 0.05 vs. siRNA Control. **e,** Immunoblotting for NRF2 protein levels in the nuclear fraction of NIH 3T3 cells treated with BDNF or vehicle (Veh). Incubation of NIH 3T3 cells with 3.7 and 7.4 nM of BDNF induces an increase in nuclear NRF2 (*N* = 3 experiments per condition). Results of statistical analyses are shown in Extended Data Fig. 5e. * *p* < 0.05 vs. vehicle (Veh). **f,** 3D imaging experiments of HT-22 cells immunofluorescence staining with BDNF and TI-VAMP (Vesicle associated membrane protein 7) antibodies. Cell nuclei were stained with DAPI. Z-stack image (MERGE z-stack) corresponds to the overlay of the three staining images, showing that the detected BDNF is observed free in the cytosol (*N* = 3 experiments). Scale bar: 10 μm.

Notwithstanding the existence of an unknown BDNF receptor or the non-specificity of the used antagonists, an alternative hypothesis to explain NRF2 activation by extracellular BDNF is that the latter enters cells by a yet to be defined mechanism. Once in the cytosol, BDNF would trigger the dissociation of KEAP1 and NRF2, allowing nuclear translocation of NRF2, as described for the intracellular p21 and p62 proteins ^11,12^. As BDNF is present cytosolic compartment (Fig. 1f), we tested whether BDNF can bind either NRF2 or KEAP1.

### BDNF interacts with KEAP1 allowing NRF2 nuclear translocation

*In silico* docking analysis of KEAP1 unveiled five potential sites of interaction with BDNF (Extended Data Fig. 2a). The two most probable docking sites included more than 44% of the best 200 poses, and involved a hydrophobic patch (Extended Data Fig. 2b and Supplementary Video 1). Surface plasmon resonance (SPR) confirmed that BDNF binds directly to KEAP1 (Fig. 2a). A binding kinetics analysis demonstrated fast association (k_a_ = 8.28 × 10^5^ M^−1^s^−1^) and low dissociation (k_d_ = 2.25 × 10^−3^ s^−1^) between BDNF and KEAP1. The equilibrium dissociation constant (K_D_ = 2.71 nM) indicated high binding affinity between BDNF and KEAP1 (Fig. 2b, righthand panel). By comparison, the affinity of phosphorylated p62, known to bind to KEAP1 and trigger dissociation from NRF2, is three orders of magnitude lower ^18^, emphasizing the exceptional high affinity of BDNF for KEAP1. Given this result, we hypothesized that BDNF and NRF2 may compete for KEAP1 binding. SPR analysis of KEAP1 on an immobilized peptide derived from the Neh2 domain of NRF2 containing an ETGE motif showed a classic interaction pattern, which was not observed when BDNF was applied (Fig. 2c). Furthermore, application of a BDNF and KEAP1 mix altered the interaction between KEAP1 and the Neh2-derived peptide (Fig. 2c). Immunoprecipitation experiments in basal conditions confirmed that BDNF binds to KEAP1 (Fig. 2d,e and Extended Data Fig. 2c). Taken together, these results indicate that, in physiological conditions, BDNF directly competes with NRF2 for KEAP1 binding. This suggests that BDNF is more than a neurotrophin acting on cognate receptors. Here we unravel a novel function for BDNF, based on a cytosolic localization and involving a direct interaction with KEAP1, which can trigger the translocation of NRF2 to the nucleus and subsequent activation of cytoprotective responses. Having outlined the mechanism of action of BDNF on the KEAP1-NRF2 system in the cytosol, we tested the other part of the hypothesis, i.e. whether extracellular BDNF can enter cells.

**Fig. 2.**
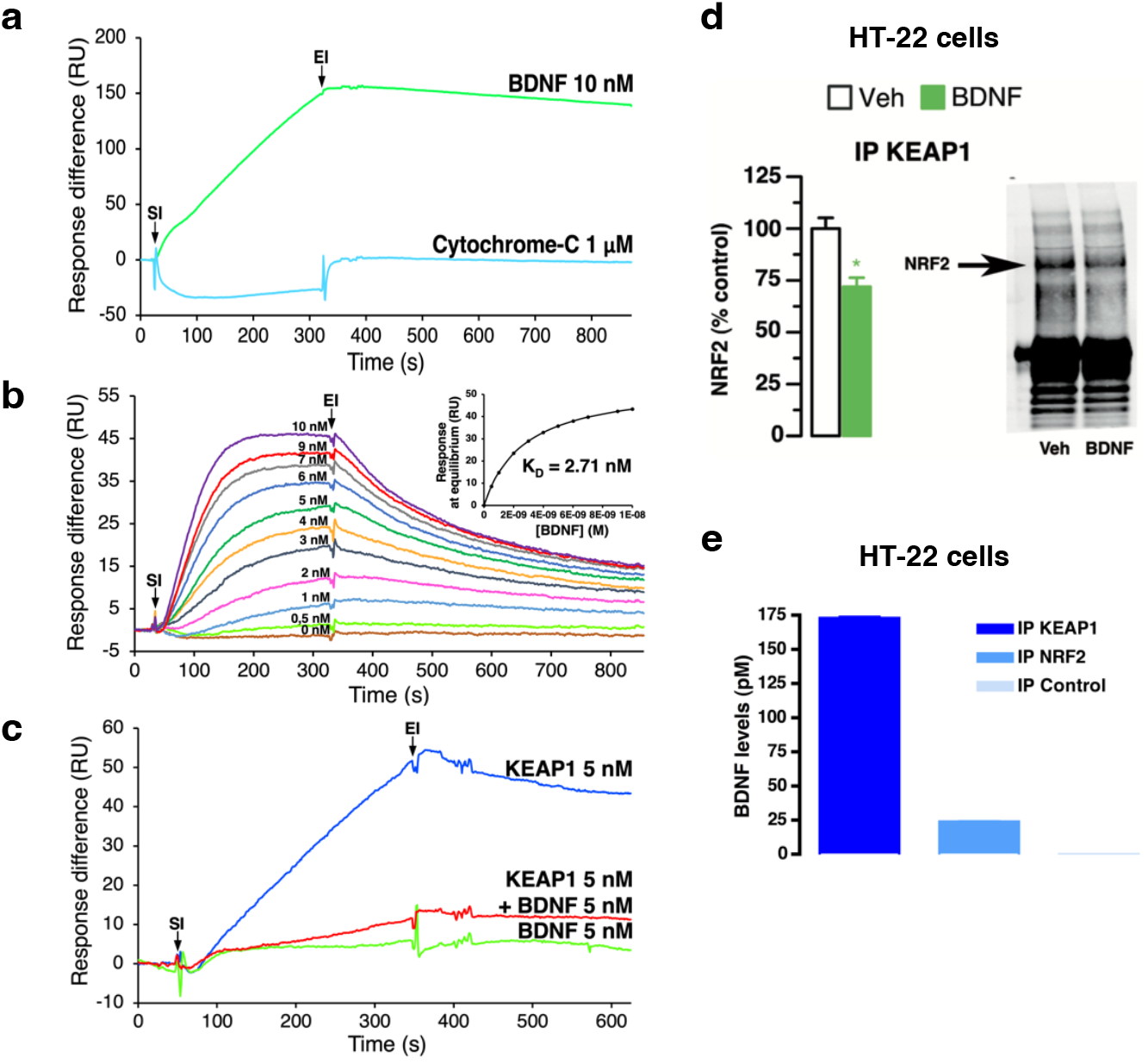
BDNF-KEAP1 cytosolic interaction. **a,** Surface plasmon resonance (SPR) was performed between KEAP1 and either BDNF (10 nM) or cytochrome C as a negative control (1 μM). SI: start of injection; EI: end of injection. Sensorgrams demonstrate a direct interaction between KEAP1 and BDNF (*N* = 3 experiments). **b,** Kinetic SPR analysis was performed between KEAP1 and increasing concentrations of BDNF (0-10 nM). Kinetic SPR analysis indicates that BDNF binds to KEAP1 with a dissociation equilibrium constant corresponding to a high binding affinity (*N* = 3 experiments). Upper panel: plot of the response at equilibrium (Req [RUs]) *versus* BDNF concentration used for the steadystate affinity fitting confirmed a KD of 2.71 nM. **c,** SPR was performed between the 16-mer ETGE domain of NRF2 and KEAP1 (5 nM), BDNF (5 nM) or a mixture of BDNF (5 nM) and KEAP1 (5 nM). Sensorgrams show a binding between the 16-mer ETGE domain of NRF2 and KEAP1, but not with BDNF, indicating that BDNF does not interact with NRF2. The BDNF/KEAP1 mixture does not interact with NRF2, indicating that BDNF intervenes in the interaction between NRF2 and KEAP1 (*N* = 3 experiments per condition). **d,** Analysis of KEAP1-NRF2 complexes by co-immunoprecipitation on cell lysates after BDNF treatment. NRF2 immunodetection performed on KEAP1 immunoprecipitates from cell lysates of HT-22 cells treated with BDNF indicates a decrease in KEAP1-NRF2 complexes in HT-22 cells incubated with BDNF (*N* = 3 experiments). * *p* < 0.05 vs. vehicle (Veh). **e,** BDNF quantification by an ELISA assay on KEAP1 or NRF2 immunoprecipitates from HT-22 cell lysates. BDNF levels in KEAP1 immunoprecipitates suggest an interaction between the two proteins. Intracellular BDNF levels are expressed per 6.0 × 10^6^ - 6.5 × 10^6^ cells (*N* = 3 experiments).

### BDNF crosses the cell membrane

To examine this question we used BDNF KO NIH 3T3 cells, which do not express BDNF in basal conditions. Addition of BDNF extracellularly resulted in a proportional accumulation of BDNF intracellularly, the latter accounting for 7.7% to 8.7% of the added extracellular concentration (Fig. 3a). As this result unequivocally demonstrates that BDNF can enter cells, we next investigated the mechanism of BDNF entry.

**Fig. 3.**
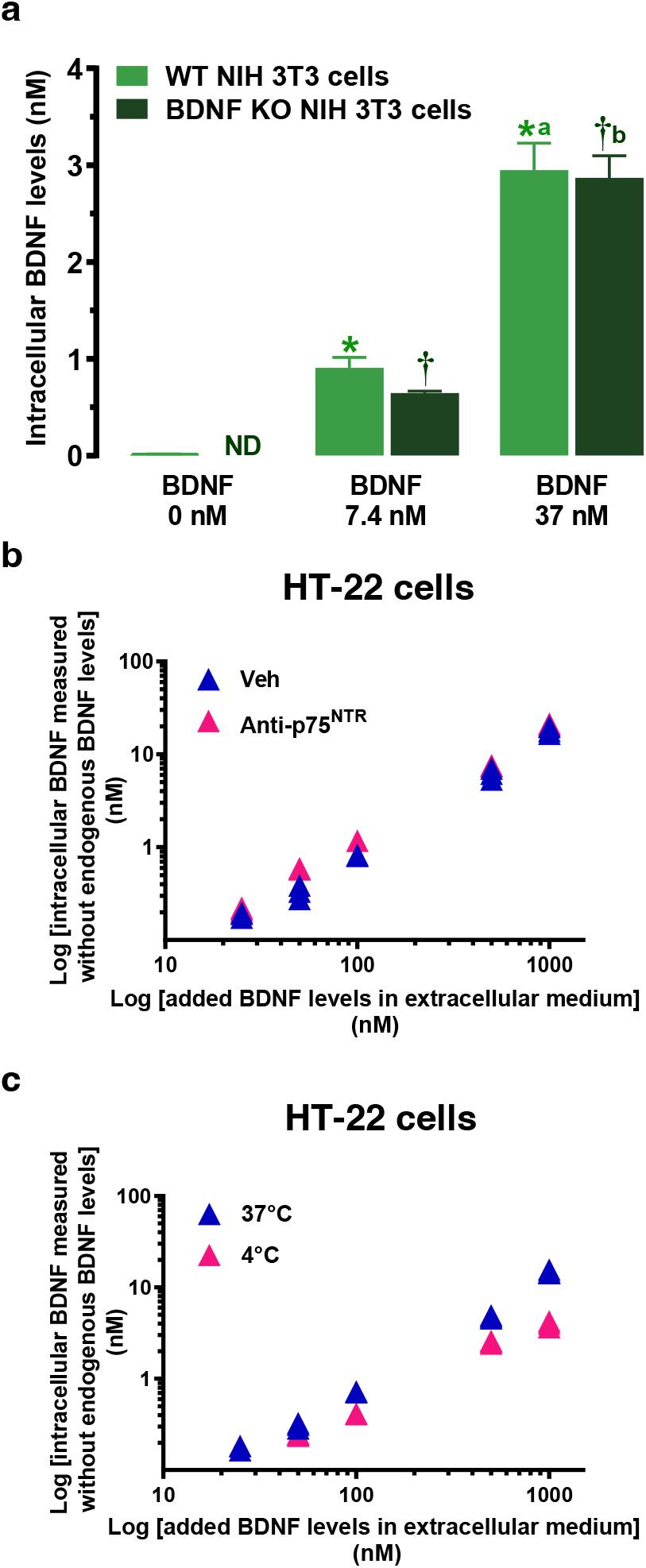
Receptor-independent internalization of BDNF. **a,** BDNF quantification on the intracellular fraction of WT NIH 3T3 and BDNF KO NIH 3T3 cells treated with BDNF (0, 7.4 or 37 nM) at 37°C. Incubation with increasing concentrations of BDNF induced an increase in intracellular BDNF levels in Wild Type (WT) NIH 3T3 and in BDNF KO NIH 3T3 cells, which do not express BDNF in basal conditions (ND: non detectable). Intracellular BDNF levels are expressed per 5.0 × 10^6^ - 5.5 × 10^6^ cells (*N* = 4 experiments). Results of statistical analyses are shown in Extended Data Fig. 5e. * *p* < 0.05 vs. WT BDNF 0 nM, † *p* < 0.05 vs. BDNF KO BDNF 0 nM, a *p* < 0.05 vs. WT BDNF 7.4 nM, b *p* < 0.05 vs. BDNF KO BDNF 7.4 nM. **b,** BDNF quantification on the intracellular fraction of HT-22 cells treated with BDNF (1.85-74 nM) and with p75^NTR^-blocking antibody or vehicle (Veh). Incubation with increasing concentrations of BDNF (1.85-74 nM) induces a linear increase in intracellular BDNF levels even in the presence of p75^NTR^ -blocking antibody. Intracellular BDNF levels are expressed per 6.0 × 10^6^ - 6.5 × 10^6^ cells (*N* = 3 experiments). **c,** BDNF quantification on the intracellular fraction of HT-22 cells treated with BDNF (1.85-74 nM) at 37°C and 4°C. Incubation with increasing concentrations of BDNF (1.85-74 nM) induces a linear increase in intracellular BDNF levels at 37°C and 4°C. Intracellular BDNF levels are expressed per 6.0 × 10^6^ - 6.5 × 10^6^ cells (*N* = 3 experiments).

We first explored the possibility of a thiol-dependent mechanism, since BDNF contains several thiol groups ^19^. This mechanism involves cell-surface protein disulfide isomerases (PDIs) ^20^. Addition of the membrane-impermeant thiol reagent DTNB (5,5’-dithiobis (2-nitrobenzoic acid)), a PDI inhibitor, did not prevent intracellular BDNF increase after extracellular BDNF supplementation, ruling out a thiol-dependent mechanism for BDNF cell entry (Extended Data Fig. 3a).

A second possibility would be cell entry *via* an active transport mechanism. If this were the case, the mechanism should saturate with increasing BDNF concentrations, and be sensitive to temperature. We found that intracellular BDNF increased along with extracellular concentration in NIH 3T3 cells in a linear fashion without saturating (Extended Data Fig. 3b). We further confirmed that the linear increase in intracellular BDNF in HT-22 cells (not altered by the antibody blocking p75^NTR^) (Fig. 3b and Extended Data Fig. 3c) correlated with the linear increase in NRF2 nuclear translocation (Fig. 1a). Importantly, the intracellular raise occurred at low temperature (4°C), nearly as efficiently as at 37 °C (Fig. 3c and Extended Data Fig. 3d). These results strongly argue against energy-dependent processes, such as endocytosis and receptor-dependent mechanisms ^21,22^.

Although the canonical view assumes that an extracellular protein exerts intracellular functions by activation of its specific receptors, our results strongly suggest that extracellular BDNF can also cross directly the cell membrane to induce a distinct intracellular effect. Specific protein-transduction domains, also known as cell-penetrating peptides (CPP), are present in a few proteins such as in the transcription factor Antennapedia or in the Trans-Activator of Transcription (Tat) protein of HIV, making them able to cross the cell membrane ^22^. CPP are short sequences (6-30 amino acid residues in length and positively charged) that confer the capacity to cross the cell membrane *via* energy-dependent and/or independent mechanisms, without the need of specific receptors ^22^. Consequently, CPP have been used as vectors for intracellular delivery of bioactive molecules ^23,24^. *In silico* analysis of the amino acid sequence of mature BDNF_(129-247)_, using CellPPD ^25^ and SkipCPP-Pred ^26^ sequence-based prediction methods, revealed the presence of a 25-amino acid sequence (223-247, KKRIGWRFIRIDTSCVCTLTIKRGR) evocative of such a protein-transduction domain at the C-terminus (C-ter BDNF). The cell penetrating properties of C-ter BDNF were investigated *in vitro* on HT-22 and NIH 3T3 cells by conjugating the peptide to *TAMRA. TAMRA-C-ter* BDNF showed the same cell membrane crossing properties at 37°C and 4°C as *TAMRA*-penetratin, a canonical CPP corresponding to the third helix of Antennapedia homeodomain (43-58) ^21,22^ (Fig. 4a, Extended Data Fig. 4a and Supplementary Video 2). In contrast, *TAMRA*-mutated C-ter BDNF peptide, predicted as non-CPP, was not internalized (Fig. 4a and Extended Data Fig. 4a). Conversely, addition of the C-ter BDNF peptide to the cell-impermeant *TAMRA* - shepherdin resulted in cell entry (Fig. 4b and Extended Data Fig 4b).

**Fig. 4.**
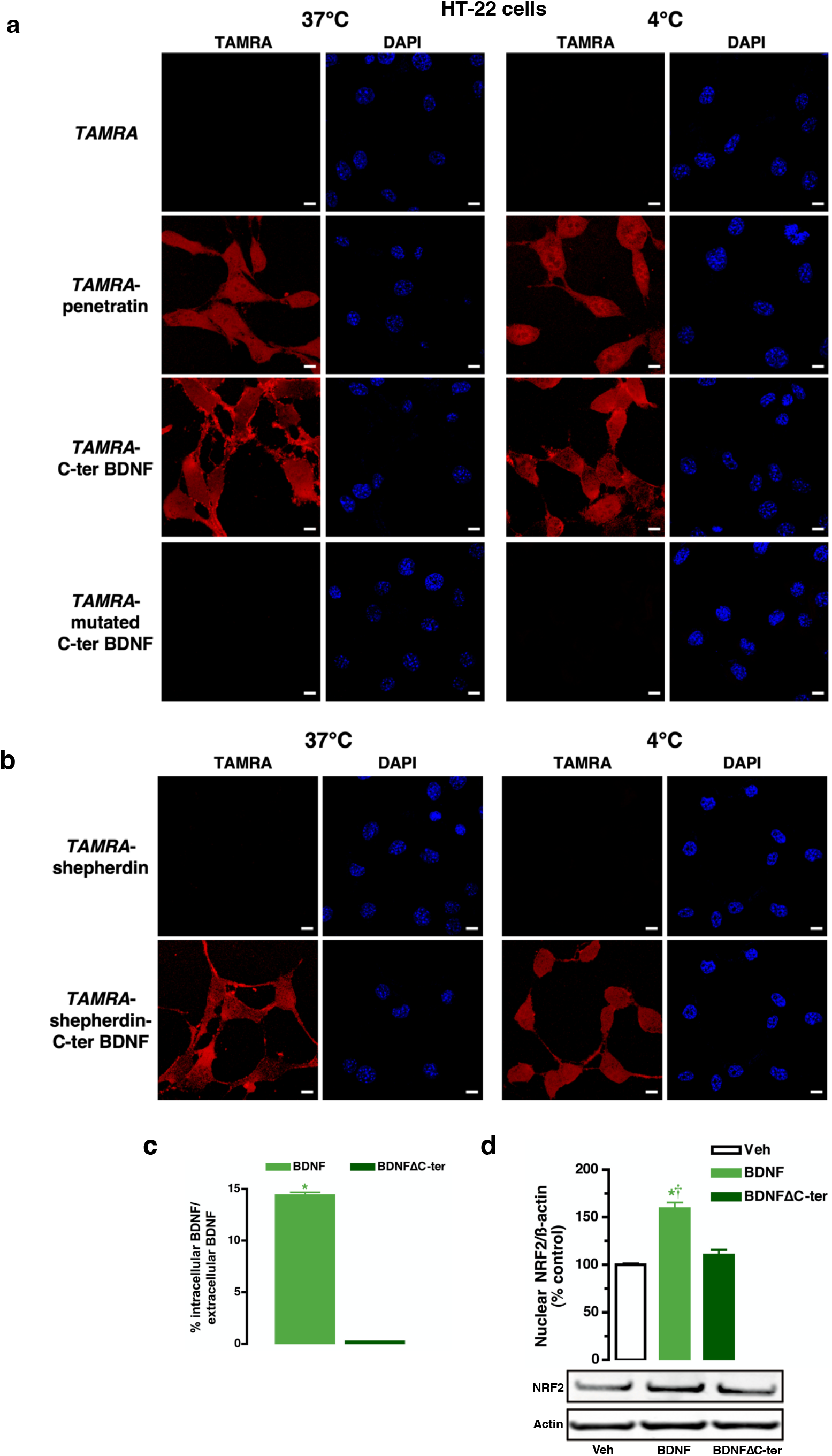
C-ter BDNF is necessary and sufficient to cross the cell membrane. **a,** Confocal microscopy of HT-22 cells incubated with *TAMRA*-labeled C-ter BDNF (1 μM), with *TAMRA*-labeled penetratin as positive control (1 μM), with *TAMRA* fluorophore (1 μM), a membrane-impermeant cargo, or with *TAMRA*-mutated C-ter BDNF (1 μM) during 20 minutes at 37°C and at 4°C. Cell nuclei were stained with DAPI. Extracellular addition of *TAMRA*-labeled C-ter BDNF to HT-22 cells resulted in its accumulation in the cytosol, like *TAMRA*-labeled penetratin. *TAMRA*-mutated C-ter BDNF does not accumulate intracellularly, showing the implication of the two mutated amino acids (*N* = 6 experiments per condition). Scale bar: 10 μm. **b,** Confocal microscopy of HT-22 cells incubated with *TAMRA*-shepherdin, an impermeant peptide, conjugated or not with C-ter BDNF (1 μM). Cell nuclei were stained with DAPI. Extracellular addition of *TAMRA*-shepherdin-C-ter BDNF to HT-22 cells results in its accumulation in the cytosol. *TAMRA*-shepherdin does not lead to cellular internalization. Each experiment has been performed at 37°C (left panels) and at 4°C (right panels); *N* = 6 experiments per condition. Scale bar: 10 μm. **c,** BDNF quantification in extracellular media and intracellular fraction of HT-22 cells after BDNF or BDNFΔC-ter addition. No intracellular BDNFΔC-ter was detected (*N* = 3 experiments per condition). * *p* < 0.05 vs. BDNFΔC-ter. **d,** Immunoblotting for NRF2 protein levels in nuclear fraction of HT-22 cells treated with BDNF or BDNFΔC-ter. Extracellular addition of BDNF to HT-22 cells, but not of BDNFΔC-ter, induces translocation of NRF2 to the nucleus (*N* = 3 experiments). Results of statistical analyses are shown in Extended Data Fig. 5e. * *p* < 0.05 vs. vehicle (Veh), † *p* < 0.05 vs. BDNFΔC-ter.

The previous experiments demonstrate that the C-ter BDNF peptide sequence is sufficient to confer cell membrane-crossing properties to otherwise impermeant molecules including peptides. To test whether the C-ter BDNF peptide sequence is necessary for BDNF to cross the cell membrane, we deleted the last 25 amino acid of the C-terminus from mature BDNF (BDNFΔC-ter). Molecular dynamics simulations indicated that BDNF conformation remains stable when this short sequence is deleted (Extended Data Fig. 4c). Supporting this proposal, SPR confirmed that BDNFΔC-ter can still bind to KEAP1 (Extended Data Fig. 4d). Furthermore, *in vivo* injection of BDNFΔC-ter into rat hippocampus led to the expected activation of p-ERK (Extended Data Fig. 4e). Thus, deletion of the CPP sequence of BDNF does not appear to alter the 3D conformation of the protein or its action on its receptors. When added extracellularly to HT-22 cells, BDNFΔC-ter was unable to enter cells (Fig. 4c) and failed to induce NRF2 translocation to the nucleus (Fig. 4d).

Taken together, these results indicate that the C-terminal sequence of BDNF is both necessary and sufficient for BDNF to cross the cell membrane. This ability was also observed in differentiated intestinal epithelial cells (Extended Data Fig. 5a,b), suggesting an ubiquitous mechanism and action, in keeping with the fact that BDNF is widely distributed in the body ^27^. Interestingly, *in silico* analysis revealed other proteins with CPP-like sequences, including numerous growth factors (Extended Data Fig. 5c). Among them, amphiregulin enters cells following a similar pattern as BDNF (Extended Data Fig. 5d).

## Discussion

The present study demonstrates for the first time that an extracellular protein known to trigger a signal transduction pathway by binding a specific receptor, can also cross the cell membrane to exert a direct action on cytosolic components. The cytosolic action of BDNF that we describe results from two remarkable properties: (i) BDNF crosses the cell membrane due to the presence of a specific amino acid sequence, and (ii) it directly interacts with KEAP1 in the cytosol to elicit NRF2 nuclear translocation and subsequent transcription of NRF2-target genes.

Our study shows that BDNF presents a functional cell-penetrating sequence that is necessary and sufficient to ensure passage through the cell membrane. Although still subject to debate, the way cell-penetrating sequences enable cell entry involves either endocytosis -as for fibroblast growth factor homologous factor 1 ^28^, which is energy-dependent- or direct translocation -which is energy-independent ^22^-. The fact that BDNF and peptides carrying its cell-penetrating sequence cross the cell membrane at low temperature in various cell types strongly argue for direct translocation. Such a translocation process may depend on the composition and physicochemical properties of the membrane. Further research is needed to determine whether alterations in those physicochemical properties, which can occur under pathological conditions ^29,30^, affect the crossing of extracellular BDNF, and ultimately its action on KEAP1-NRF2.

The known proteins capable of activating NRF2 by directly interacting with KEAP1 and disrupting the KEAP1-NRF2 complex, generally contain a canonical or non-canonical ETGE motif and belong to the category of intracellular proteins without exception ^10^. Hence they represent functional intermediaries linking some intracellular processes to the KEAP1-NRF2 cytoprotective system. BDNF lacks these two characteristics: on the one side, its ability to interact with KEAP1 does not rely on an ETGE motif, yet it might involve hydrophobic patches; on the other side, it originates from the extracellular environment. The fact that the affinity of BDNF for KEAP1 is three orders of magnitude higher than that of p62 -an ETGE-containing disruptor-indicates that BDNF could be one of the main regulators of KEAP1-NRF2 once in the cytosol. As, according to our results, cytosolic BDNF levels are directly dependent upon extracellular concentration, we propose that extracellular BDNF could act as a rheostat, continuously controlling the extent of NRF2 activation in cells. This mechanism would be complementary to the well-known receptor-dependent effects exerted after its activity-dependent release from neurons ^1,2^.

The cytoprotective action of BDNF can rely not only on the activation of its receptors and their signal transduction pathways ^31^, but also on its cytosolic action on the KEAP1-NRF2 cytoprotective system. As this receptor-independent pathway occurs *in vivo*, the functional link between BDNF and NRF2 opens new insights into diseases where antioxidant and neuroprotective functions are essential, including cancer, neurodegenerative diseases and stress-related disorders ^5,6,32^. BDNF levels are often decreased in pathological conditions, which would result in a diminished translocation of NRF2 to the nucleus and a less efficient activation of defense mechanisms as observed in vulnerability to depression ^5^. When BDNF levels raised, as during the recovery of resilient animals, expression and functionality of NRF2-dependent anti-oxidant enzymes, such as superoxide dismutase and the sulfiredoxin/peroxiredoxin system, were restored ^5^. The expression of these anti-oxidant enzymes is also activated by BDNF in cell culture. By discovering an unsuspected mode of action of BDNF, serving as a predictor of a depression profile, we propose a new therapeutic target in diseases presenting an alteration of both BDNF levels and KEAP1-NRF2 cytoprotective system.

The fact that other extracellular proteins, including several growth factors, contain CPP-like sequences, underlines the key role of this novel signaling pathway for the receptor ligand proteins which is an energy-independent process having a low cost for cell metabolism. An unavoidable step will be to determine their intracellular actions after cell entry. Besides the canonical autocrine and paracrine signaling for growth factors, this novel pathway of cell entry to exert an intracellular function opens exciting new avenues in cell signaling and a new research agenda in cell biology.

## Methods

### Reagents

All experiments were performed using the BDNF mature recombinant (GF029; Merck Millipore, USA) or the C-ter truncated form of BDNF (BDNFΔC-ter), originated from a purified mature form of human BDNF as described below. Preliminary experiments showed no difference on cellular uptake and nuclear NRF2 translocation in these two human mature BDNF. The mature form of human BDNF (NCBI GI 987872, Residues 128-247) and the truncated form deleted from the C-terminal CPP sequence (BDNFΔC-ter;Residues 128-222) were expressed in *E. coli* using the pET28a vector (GenScript, USA). BDNF was diluted either in saline for *in vivo* experiments or in DMEM for *in vitro* experiments. The blocking receptor antibody was a rabbit polyclonal antiserum to the extracellular domain of mouse p75^NTR^ (9650), provided by M.V. Chao, diluted to 1:100 ^33^. ANA12 (10 μM [cells] or 1 μg [rats] ^34,35^; SML0209, Sigma-Aldrich/Merck group, USA) was dissolved in DMSO / Tween 80 / sterile 0.9% NaCl solution. Quercetin (10^−5^ M, 00200595; Sigma-Aldrich/ Merck group, USA) ^36^, amphiregulin (10 or 50 nM, ab229600; Abcam, Netherlands) and DTNB (2.5 mM, D8130; Sigma-Aldrich/Merck group, France) was diluted in DMEM. BDNF concentrations in different cell fractions were determined by using a BDNF ELISA kit (KE00096; detection range: 12.5 – 800 pg/mL; Proteintech Europe, United Kingdom). Amphiregulin concentrations in HT-22 cell fractions were determined by using a Human Amphiregulin ELISA kit (ab222504; detection range: 18 – 600 pg/mL; Abcam, Netherlands). Protein concentrations were measured using RC DC Protein Assay (5000120; Bio-Rad, USA). Surface Plasmon Resonance (SPR) experiments were performed using KEAP1 (11981-HNCB; Sino Biological Inc., China) and 16-mer ETGE NRF2 peptide synthetized by the Molecular Interaction Platform (Institut de Biologie Paris Seine, France). *TAMRA*(5-Carboxytetramethylrhodamine)-peptides were synthetized by ProteoGenix (France).

### Animals

Male Sprague-Dawley rats weighing 290-310 g (9 weeks old at the beginning of the experiments) were obtained from the same breeder (Centre d’élevage R. Janvier, 53940 Le Genest-St-Isle, France). On their arrival at the laboratory, they were housed together for 10 days before micro-infusion experiments in a chronobiological animal facility (Enceinte Autonome d’Animalerie, A110SP; Thermo Electron Corporation, Saint Herblain, France) under controlled environmental conditions (21 ± 1 °C, 60% relative humidity, 12/12 hr light-dark cycle with lights on at 7 am, food and water *ad libitum*). Procedures involving the animals and their care were performed in accordance with institutional guidelines, which complied with national and international laws and policies (French Government directive #87-848, October 19, 1987; authorization #R-75ParisV-F1-12-rongeurs to Chrystel Becker (project number: 2019022115289223).

### *In vivo* microinfusion

Microinjection of BDNF, BDNFΔC-ter, ANA12 and p75^NTR^-blocking antibody (anti-p75^NTR^) was performed on the rats after they had been anesthetized with ketamine (33 mg/kg; Clorkétam 1000; Vétoquinol, France) and xylazine (6 mg/kg; Rompun 2%; Bayer, Germany) and placed in a stereotactic apparatus (Kopf Instruments; Phymep, France). Stainless-steel cannulae (Plastic One; Phymep, France) were implanted bilaterally to the dorsal hippocampus at the following stereotactic coordinates with respect to the bregma: - 4.8 mm anteroposterior, ±3.2 mm mediolateral, and −2.2 mm dorsoventral (from dura). The cannulae were anchored to the skull using acrylic dental cement. A 15-min pre-infusion of 1 μg of ANA12 ^35^ and anti-p75^NTR^ (antiserum 9650, 1:100) ^33^ or of vehicle was followed by infusion of 1 μg BDNF, BDNFΔC-ter or ANA12, anti-p75^NTR^ and BDNF, or of vehicle (Veh). 50 min after, animal brains were removed, and the hippocampal injected areas dissected and subjected to subcellular fractionation, then to western blot analysis.

### Cells

Hippocampal HT-22 cells, BDNF KO NIH 3T3 cells (CRISPR-U™ technology; Ubigene, China), NIH 3T3 cells (WT NIH 3T3 cells, Ubigene, China) and intestinal differentiated Caco-2 cells were cultured and maintained at 37 °C under an atmosphere containing 5% CO_2_ in Dulbecco’s Modified Eagle Medium (DMEM; 41966029; Gibco/Thermo Fisher Scientific, USA) supplemented with 5-10% fetal bovine serum (F7524; Sigma-Aldrich/Merck group, USA) and 1% penicillin/streptomycin (15140122; Gibco/Thermo Fisher Scientific, USA) (HT-22 and Caco-2 cells) or in ATCC-formulated Dulbecco’s Modified Eagle’s Medium (DMEM; ATCC-30-2002; ATCC, USA) with 10% bovine calf serum (ATCC-30-2030; ATCC, USA) and 1% penicillin/streptomycin (WT and BDNF KO NIH 3T3 cells). Being unable to have the necessary control showing that the labeling of BDNF would not alter the transmembrane passage of the protein, we used BDNF KO NIH 3T3 cells. Cells were incubated with BDNF, BDNFΔC-ter, *TAMRA*-peptides, quercetin (Querc.,10 μM), amphiregulin or vehicle (Veh) in DMEM (without serum) for 20 min at 4°C or for 40 min at 37°C. The cells were then subjected to subcellular fractionation to obtain nuclear and cytosolic extracts. For the experiments at 4 °C, the cells were placed at 4 °C for 20 min. For experiments requiring receptor blockade, HT-22 cells were pre-incubated with the p75^NTR^-blocking antibody (antiserum 9650, 1:100) ^33^ or with vehicle for 15 min and then incubated with p75^NTR^-blocking antibody and BDNF or vehicle. Caco-2 cells, which express TrkB and p75^NTR^ receptors ^37,38^ were pre-incubated with ANA12 (10 μM) ^34^ and p75^NTR^-blocking antibody for 15 min and then incubated with ANA12 + p75^NTR^-blocking antibody and BDNF or vehicle. For medium enriched with different doses of BDNF, the extracellular and intracellular BDNF concentrations were determined by using a BDNF ELISA kit (KE00096; Proteintech Europe, United Kingdom). Results were expressed either as log[extracellular BDNF levels] *versus* log[intracellular BDNF levels (without endogenous BDNF for HT-22 cells)], or as BDNF levels (pM or nM) per 6.0 × 10^6^ - 6.5 × 10^6^ cells for HT-22 cells, per 2.5 × 10^5^ cells for Caco-2 cells or 5.0 × 10^6^ – 5.5 × 10^6^ cells for NIH 3T3 cells. Endogenous BDNF values were 12.54 ± 1.03 pM (HT-22 cells). For the BDNFΔC-ter experiments, when 37 nM of BDNF was added to the extracellular medium, 5.32 nM of BDNF-like immunoreactivity was measured in the cells (*i.e*. 14% of the total immunoreactivity present in the extracellular medium). The crossreactivity determined for BDNFΔC-ter was 3%. When 111 nM of BDNFΔC-ter was added, we detected 33.3 nM of extracellular BDNF-like immunoreactivity and 0.02 nM of intracellular BDNF-like immunoreactivity (*i.e*. 0.07% of the total immunoreactivity present in the extracellular medium). Experiments with Poly(sodium) 4-styrenesulfate (PssNa) (243051; Sigma-Aldrich/Merck group, USA), a polyelectrolyte able to scavenge charged molecule ^39^, excluded that BDNF measured in the intracellular fraction corresponds to a BDNF bound to the extracellular part of the plasma membrane.

### siRNA knockdown

BDNF knockdown was performed using either Silencer® Select siRNA targeting BDNF or a nontargeting Silencer® Select siRNA as a control (Thermo Fisher Scientific, USA). siRNA transfections were performed at a final concentration of 50 nM, using Lipofectamine RNAiMAX (Thermo Fisher Scientific, USA) according to the manufacturer’s instructions. BDNF silencing with siRNA had an efficiency of 82%: 17.62 ± 0.52 pg/mg protein for siRNA control versus 3.13 ± 0.17 pg/mg protein for siRNA BDNF.

### Nuclear and cytosolic extracts

After scrapping in PBS with protease inhibitor cocktail, cells were centrifuged (500x *g*, 5 min, 4 °C). After removing the supernatant, cells were suspended in ice-cold buffer A (10 mM HEPES pH 7.5, 150 mM NaCl, 1 mM EDTA, protease inhibitor cocktail [5 μl/ml] and phosphatase inhibitor cocktail [10 μl/ml]) and 0.5% Nonidet P-40) (I8896; Sigma-Aldrich/Merck group, USA). After 10 min on ice, samples were centrifuged (1200x *g*, 10 min, 4 °C). Supernatants were saved as the cytosolic fraction. Pellets were washed in ice-cold buffer A and sedimented by centrifugation at 1200 x *g* for 10 min at 4 °C. Nuclei were resuspended in ice-cold buffer B (20 mM Hepes pH 7.5, 420 mM NaCl, 1.2 mM MgCl2, 0.2mM EDTA, 25% glycerol, 0.5mM DTT, protease inhibitor cocktail [5μl/ml] and phosphatase inhibitor cocktail [10 μl/ml]). Samples were sonicated three times (25 s, 4 °C) every 10 min. The cytosolic and nuclear fractions were centrifuged at 20,000 x *g*, for 30 min at 4 °C. Supernatants were rapidly frozen in liquid nitrogen and stored at −80 °C. A similar protocol was used for hippocampal extracts after Dounce homogenization in ice-cold buffer A^6^.

### Western blotting

20 μg (cultured cells) or 30 μg (hippocampal) of nuclear or 40 μg of cytosolic proteins were resolved by SDS-polyacrylamide gel electrophoresis in a 12% polyacrylamide gel (NuPAGE Gel, NP0335BOX; Thermo Fisher Scientific, USA). Proteins were transferred to nitrocellulose membranes (Thermo Fisher Scientific, USA) for immunoblotting according to standard procedures. Antibody dilutions were as follows: 1:250 for rabbit polyclonal antibodies against NRF2 (sc-13032; Santa Cruz Biotechnology, USA), 1:500 for rabbit polyclonal antibodies against KEAP1 (sc-15246; Santa Cruz Biotechnology, USA), erk-p (clone AW39R, 1/500, Merck Millipore, USA), p75 antibody (1/1000, 9992), and 1:10,000 for the fluorescent IRDye® 800CW secondary antibody (Eurobio Ingen, France). Membranes were also probed with either a mouse monoclonal antibody against GAPDH or a mouse monoclonal antibody against beta-actin (dilutions of 1:10,000 and 1:1000; Sigma-Aldrich/Merck group, USA) and with the IRDye® 680CW secondary antibody from Eurobio Ingen (926-32220; 1:10,000), in order to correct for protein loading. Images were acquired with the Odyssey® infrared imaging system (LI-COR Biosciences, USA). Protein band intensities were quantified with Odyssey® software (version 3.0; LI-COR Biosciences, USA).

### Immunofluorescence

Cells were cultured on polylysine coverslips. Cells were incubated with BDNF (7.4 or 37 nM) or vehicle for 40 minutes at 37°C. After three washes in PBS, cells were fixed with 2.5% paraformaldehyde for 20 min at room temperature. Permeabilization was performed in PBS / 0.5% Triton X-100 for 5 min. Cells were incubated with BDNF (1/50, sc-546; Santa Cruz Biotechnology, USA), TI-VAMP (clone158.2, 1/500) antibodies overnight at 4 °C, washed three times with PBS, incubated with secondary antibodies (1/100) for 1 hr at room temperature, and then with WGA Alexa Fluor 647 conjugate (1/500) for 10 min. Cell nuclei were stained with DAPI. Coverslips were mounted with ProLong (P36930; Invitrogen/Thermo Fisher Scientific, USA) and analyzed by confocal microscopy. Immunofluorescence images were captured with a Leica SP2 confocal laser (Wetzlar, Germany). Images were analyzed using Leica Confocal Software, and mounted using Image J and Photoshop CS6. The median sections (*i.e*. Z = 005) are displayed here. 3D imaging experiments were performed by using Leica SP8 3D STED of NeurImag Platform (IPNP; Paris, France). LAS software (Leica) was used to acquire and analyze images.

### Cellular uptake of *TAMRA*-labeled peptides

Cells were cultured on polylysine coverslips and incubated with the *TAMRA*-C-ter BDNF (*TAMRA*-KKRIGWRFIRIDTSCVCTLTIKRGR-COOH; Proteogenix, France) corresponding to the 25 amino acids of the C-terminal sequence of BDNF at 1 μM, with 1 μM mutated *TAMRA*-C-ter BDNF peptide (CellPDD predicting score = 0.03, *i.e*. mutation abolishing the CPP characteristics : *TAMRA*-KKEIGWRFIRIDTSCVCTLTIKEGR-COOH; Proteogenix, France), with 1 μM *TAMRA*-penetratin (positive control, *TAMRA* -RQIKIWFQNRRMKWKK-COOH; Proteogenix, France), with 1 μM *TAMRA*-sheperdin (*TAMRA*-KHSSGCAFL-COOH; Proteogenix, France), with 1 μM *TAMRA*-sheperdin-C-ter BDNF (*TAMRA*-KHSSGCAFLKKRIGWRFIRIDTSCVCTLTIKRGR-COOH; Proteogenix, France), or with *TAMRA* fluorophore alone for 20 minutes. *TAMRA* is a well-admitted membrane-impermeant cargo to test the transmembrane passage of CPPs ^40^. After washing, cells were fixed with 4% paraformaldehyde for 10 min at room temperature. After washing, cell nuclei were stained with DAPI (4’,6-diamindino-2-phenylindole). The coverslips were mounted with ProLong (P36930; Invitrogen/Thermo Fisher Scientific, USA) and analyzed by confocal microscopy (Zeiss LSM710, 93x objective). ZEN software was used to acquire and analyze images.

### Fluorescence videomicroscopy

Cells were cultured in a 12 well-plate and incubated with 1 μM *TAMRA*-C-ter BDNF (*TAMRA*-KKRIGWRFIRIDTSCVCTLTIKRGR-COOH; Proteogenix, France) for 40 minutes in incubation chamber at 37°C in a 5% CO_2_ atmosphere. Acquisition of live cells was performed every 30 seconds for 40 minutes. After washing, fluorescence and Differential Interference Contrast (DIC) images were also taken. Images were taken using a CMOS camera (ORCA Flash 2.8; Hamamatsu Photonics) mounted on an epifluorescence microscope system (AxioObserver Z1; ZEISS, 40x objective). ZEN software was used to acquire and analyze images.

### Immunoprecipitation

HT-22 cells were treated for 40 min with either BDNF or saline as a control. After treatment, they were washed three times with cold PBS, incubated for 5 min in lysis buffer containing 50 mM Tris-HCl (pH 7.5), 150 mM NaCl, 1% Nonidet P-40, and phosphatase and proteinase inhibitor cocktails (P5726 and P2714; Sigma-Aldrich/Merck group, USA), and scrapped. The homogenates were centrifuged at 10,000 *g* for 10 min at 4 °C. Cell lysates (1 mg of proteins) were incubated with 2 μg of anti-KEAP1 antibodies for 30 min in lysis buffer, and 100 μL of μMACSTM Protein A/G MicroBeads (Miltenyi Biotec, Germany) were added for 1 hr incubation before applying to pre-equilibrated μ columns. Columns were washed five times with wash buffer. Elution was performed by applying 20 μL of pre-heated (95 °C) SDS buffer (50 mM Tris-HCl pH 6.8, 10% glycerol, 35 mM SDS and 50 mM DTT) to the columns. Immunoprecipitated proteins after KEAP1 immunoprecipitation were subjected to NRF2 western blot analysis. To assess the interaction between BDNF and either NRF2 or KEAP1, cells were lysed with RIPA buffer containing 50 mM Tris-HCl (pH 7.5), 150 mM NaCl, 0.5% Nonidet P-40, and phosphatase and proteinase inhibitor cocktails. The lysates were treated as described above, except for the ultrasonic bath. Immunoprecipitations with KEAP1, NRF2 and control protein were performed as described above. The immunoprecipitated fractions for the vehicle conditions served to determine BDNF levels using an ELISA kit (KE00096; Proteintech Europe, United Kingdom).

### Molecular modeling

Protein-protein docking was performed in the Discovery Studio 2018 Modeling Environment (Release 2018; Dassault Systèmes, Vélizy-Villacoublay, France) using the ZDOCK protocol ^41^. This protocol computed a rigid body docking of BDNF (as ligand) on KEAP1 (as receptor), outputting 2000 poses. The poses were then re-ranked with the ZRANK function, which has been shown to be more accurate ^42^. To analyze the results, poses were clustered according to an all-against-all root mean square deviation (RMSD) matrix. For RMSD calculation, only the heavy atoms of the interface residues of a pose were considered (according to a distance cutoff of 10 Å from the receptor), so the matrix was nonsymmetrical ^43^. Based on an RMSD cutoff of 10 Å, a cluster center was selected (pose with the most neighbors) and grouped with its neighbors as a cluster before being removed from the matrix in order to process the next cluster. Clustering stopped when 100 clusters had been found or when no more poses were available to act as cluster centers. We then eliminated all non-clustered poses. For docking region analysis, we displayed the top 100 poses belonging to the 10 largest clusters. We defined regions by grouping all the poses that were at less than 10 Å from their neighbor. For each region, we selected the cluster centers and enlarged the selection to every cluster for which a pose was within a radius of 10 Å. This defined the clusters belonging to each region. Molecular dynamics simulation was performed in the Discovery Studio Modeling Environment (Release 2018; Dassault Systèmes, Vélizy-Villacoublay, France) using CHARMm 42.2 ^44^ and NAMD 2.13 ^45^. The protein was placed in an orthorhombic box at a minimum distance of 10 Å from the boundaries and solvated with a pre-equilibrated solvent containing TIP3 waters and Na^+^/Cl^−^ counterions at a concentration of 0.15 M. The system was then minimized, thermalized and equilibrated at 300 K for production. Finally, a 50-ns production run was performed in the NPT ensemble at 300 K, using a Langevin thermostat and piston, for temperature and pressure control, respectively. The RMSD analysis was done in the Discovery Studio.

### Surface Plasmon Resonance (SPR)

Binding experiments and kinetics assays were carried out using a BIACORE 3000 instrument (GE Healthcare, USA). Two different sensor chips were used, namely a CM5 sensor chip for primary amine group immobilization and a streptavidin sensor chip (SA) for biotin immobilization. Binding and kinetic experiments were carried out at a flow rate of 5 μl/min, with 5-min contact time, using HBS-EP (10 mM HEPES, pH 7.4, 150 mM NaCl, 3 mM EDTA, 0.005% surfactant P-20) as running buffer. SPR experiments on CM5 sensor chip: After sensor surface activation by injection of a mixture of 0.2 M 1-ethyl-3-(3-dimethylaminopropyl) carbodimide hydrochloride (EDC) and 0.05 M *N*-hydroxysuccinimide (NHS) (flow rate: 10 μl/min, contact time: 7 min), KEAP1 (11981-HNCB; Sino Biological Inc., China) was covalently immobilized through primary amino groups to the carboxy-methylated dextran matrix of CM5 sensor chip. A solution of 100 nM KEAP1 diluted in immobilization buffer (10 mM sodium acetate pH: 4.5) was injected at a flow rate of 10 μl/min during 7 min. The immobilization was followed by injection of 70 μl of 1 M ethanolamine hydrochloride, pH 8.5, at a flow rate of 10 μl/min to saturate the free activated sites of the carboxy-methylated dextran matrix. A reference surface without protein was prepared using the same procedure. The analytes BDNF, BDNFΔC-ter or Cytochrome-C (negative control which has a molecular weight and a pHi similar to those of BDNF; C2037; Sigma-Aldrich/Merck group) were injected on immobilized KEAP1. Kinetic sensorgrams were obtained by passing various concentrations of the analyte (BDNF) over the ligand with a 5-min association phase and an 8-min dissociation phase. The sensor surface was regenerated between each cycle (association - dissociation) with one injection of 10 mM Glycine hydrochloride (pH 1.5) at a flow rate of 30 μl/min during 30 sec. Identical injections over blank surface areas run in parallel (and giving a value of 0 Response Unit [RU]) were subtracted from all experiments. Sensorgrams were analyzed using BIAevaluation 4.1 software (GE Healthcare, USA). Kinetic fittings were performed using the Langmuir 1:1 interaction model embedded in the BIAevaluation software. The quality of the fit was assessed by the statistical chi2 value provided by the software (chi2 values < 10 were considered acceptable). SPR experiments on SA sensor chip: A biotinylated peptide containing the *ETGE* motif of NRF2 with a 3 x glycine spacer (Biotin-GGG-69AFFAQLQLDEETGEFL84-COOH) was synthetized by a peptide synthesis platform (IBPS-FR 3631; Sorbonne University, Paris, France), using Fmoc chemistry and a Liberty Blue automated microwave peptide synthesizer (CEM Corporation, Matthews, NC, USA). The peptide was diluted to 50 nM in running buffer (HBS-EP) and immobilized on a streptavidin sensor chip (sensor chip SA, GE Healthcare, USA). The analytes KEAP1, BDNF or a BDNF/KEAP1 mixture (25 min incubation at room temperature) were injected on the immobilized NRF2 peptide. The sensor surface area was regenerated between each cycle (association-dissociation) by two successive injections of 40 mM OGP and 1M NaCl/50 mM NaOH at a flow rate of 30 μL/min and 30 sec contact time. Identical injections over blank surface areas run in parallel (and giving a value of 0 RU) were subtracted from all experiments. Sensorgrams were analyzed as described above.

### Real-time-qPCR

*\* RNA extraction*: Total RNA was isolated from frozen HT-22 cells or brain tissue with the RNeasy Mini Kit (Qiagen, Courtaboeuf, France). Total RNA from living HT-22 cells were isolated using Trizol (Thermo Fisher Scientific, USA). RNA quality and concentration were evaluated with a NanoDrop spectrometer (Thermo Fisher Scientific, USA). *\* RT-qPCR*. For real-time PCR analysis, first-strand cDNA synthesis (0.6 μg of total RNA per 15 μl reaction) was performed using olig-dT (Reverse Transcription Core Kit; Eurogentec, Seraing, Belgium). Real-time PCR was carried out with the Light Cycler Fast Start DNA Master SYBR Green Kit (Roche Applied Science, Germany) using LightCycler (Roche Applied Science, Germany) and primers for target mRNA, namely: Ntrk2 (NM_001025074) and Ppia (CyCA, NM_012583) provided by Eurogentec (Seraing, Belgium). The difference in mean threshold cycle (Δ Ct) values was determined using the second derivative maximum method. To check for the presence of Ntrk2 in brain tissue and HT-22 cells, PPIA was used as a positive control, and results were expressed in absolute values (Ct). * *Custom TaqMan Array Cards for NRF2 target genes*. After quantification and integrity evaluation of samples, retrotranscription was performed. TaqMan low-density array (TLDA) cards (Thermo Fisher Scientific, USA) were used to determine the transcriptional expression of 90 NRF2 target genes and 4 reference gene candidates (*GAPDH, PPIA, B2M, GUSB*). Two reference genes (*GAPDH, GUSB*) were selected for their stability. Expression of genes was analyzed by the comparative 2^−ΔCq^ method with ΔCq = Cq (target gene) – Cq (mean of the housekeeping genes). Relative expression of target genes was quantified using the 2^−ΔΔCq^ method.

### Statistical analyses

Differences in nuclear NRF2, cytosolic NRF2, cytosolic KEAP1, intracellular BDNF levels, were subjected to one-way analyses of variance (ANOVAs). Group effects and treatment effect were subjected to two-way analyses of variance. All the results yielded by the ANOVAs (*F* values and *p* values) are provided in Extended Data Fig. 5e. If an ANOVA revealed a significant effect, we ran a *post hoc* Bonferroni test to determine whether differences were truly statistically significant. All data are presented as the mean ± standard error of the mean (*SEM*). Non-parametric Mann Whitney U test was used to assess differences between the groups for NRF2 target genes expression.

## Data availability

The datasets generated during the current study are available from the corresponding author on reasonable request (Chrystel Becker).

## Acknowledgments

This research was supported by grants from Institut National de la Santé et de la Recherche Médicale (INSERM), Sorbonne Université, Fédération pour la Recherche sur le Cerveau (FRC), and Agence Nationale de la Recherche (ANR). Hippocampal HT-22 cells were generously provided by Dr David Schubert (The Salk Institute, San Diego, CA, USA), anti-p75^NTR^ antibodies by Prof. Moses Chao (The Salk Institute, San Diego, CA, USA), and TI-VAMP antibodies (clone158.2) by Dr Lydia Danglot (IPNP, Paris, France). We acknowledge NeurImag facility of the Institute of Psychiatry and Neuroscience of Paris where 3D imaging experiments were carried out. We also acknowledge Lionel Forestier from BISCEm (Biologie Intégrative Santé Chimie Environnement) platform (Université de Limoges, France) for carrying out the TaqMan Gene Expression Assays. We thank Sylvie Riquier for their invaluable advice. The authors would like to thank Prof. Mario Ollero for his critical reading of the manuscript and helpful suggestions.

## Author contributions

J.F. performed experiments, analyzed data, interpreted data and contributed to writing the manuscript; F.B. designed research, performed experiments, analyzed data, interpreted data and wrote the manuscript; A.C performed molecular modeling and contributed to writing the manuscript; D.C. performed RT-PCR experiments and participated to *in vivo* microinjections and *in vitro* experiments; P.N. interpreted data, advised and contributed to writing the manuscript; V.P. participated to *in vitro* experiments; S.C. participated to *in vitro* experiments; T.B. supervised SPR experiments; E.B. participated to *in vitro* experiments; M.S. participated to BDNF and BDNFΔC-ter production; J.M. participated to *in vitro* experiments; A.P.F. participated to immunocytochemistry; P.C. contributed to writing the manuscript; F.A. supervised molecular modeling; M-P.G. performed and supervised BDNF and BDNFΔC-ter production; G.C. interpreted data and designed research; G.T. performed *in vitro* experiments, interpreted data and designed research; C.B. designed research, analyzed data, interpreted data, advised and supervised the project and wrote the manuscript. J.J.B. designed research, analyzed data, interpreted data, advised and supervised the project and wrote the manuscript. C.B. designed research, performed experiments, analyzed data, interpreted data, advised and supervised the project and wrote the manuscript.

## Additional information

**Extended data** is available for this paper.

**Supplementary Information** is available for this paper: Supplementary Videos 1,2.

**Correspondence and requests for materials** should be addressed to Chrystel Becker.

