## Supplementary Data for "A receptor-independent signaling pathway for BDNF"

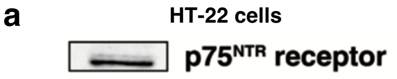


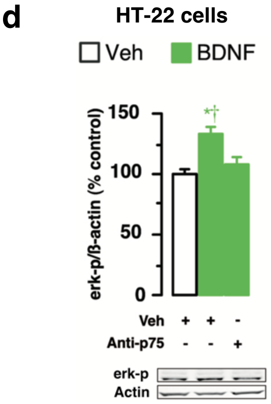


b

| Brain tissue | | HT-22 cells | |
|---|---|---|---|
| Ppia | Ntrk2 | Ppia | Ntrk2 |
| 21.17 | 30.04 | 20.46 | ND |

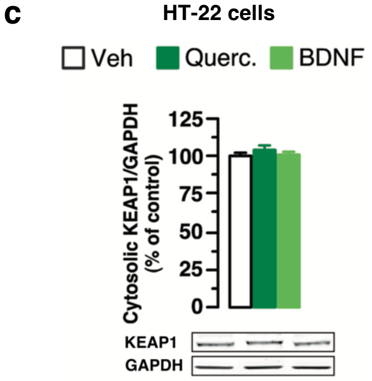


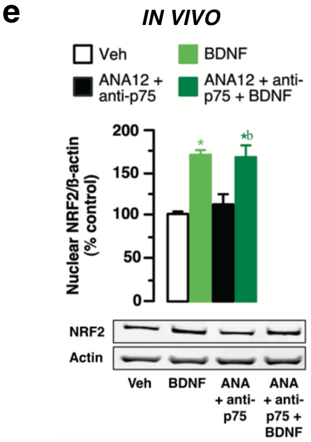


**Extended Data Fig. 1 | BDNF controls NRF2 nuclear translocation in a receptor-independent manner. a,** Immunoblotting for $p75^{NTR}$ receptors in HT-22 cells. HT-22 hippocampal cells express constitutively $p75^{NTR}$ receptors. **b,** Ntrk2 mRNA detection in brain tissue and HT-22 hippocampal cells. Ppia is used as positive control. Ntrk2 mRNA is not detected in HT-22 hippocampal cells, confirming that these cells lack TrkB receptors, in contrast to brain tissue. Results are expressed in absolute values (Ct). ND = not detectable. **c,** Immunoblotting for KEAP1 protein levels in the cytosolic fraction of HT-22 cells treated with BDNF (7.4 nM) or quercetin (Querc., 10 μM) as positive control. Incubation of HT-22 cells with BDNF or quercetin has no effect on cytosolic KEAP1 protein levels ($N$ = 3 experiments). Results of statistical analyses are shown in Extended Data Fig. 5e. **d,** Immunoblotting for phospho ERK (erk-p) in HT-22 cells treated with BDNF or vehicle (Veh). Addition of BDNF to HT-22 cells leads to activation of the ERK pathway. Blockade of $p75^{NTR}$ receptors with an anti- $p75^{NTR}$ antibody, followed by addition of BDNF, prevents activation of the ERK pathway ($N$ = 3 experiments). Results of statistical analyses are shown in Extended Data Fig. 5e. * $p < 0.05$ vs. vehicle (Veh), † $p < 0.05$ vs. BDNF + Anti-$p75^{NTR}$. **e,** Immunoblotting for NRF2 protein levels in nuclear extract of rat hippocampus after *in vivo* infusion of BDNF (1 μg) or vehicle (Veh). BDNF increases the protein levels of nuclear NRF2. Blockade of TrkB and $p75^{NTR}$ receptors with the TrkB inhibitor ANA12 and an anti- $p75^{NTR}$ antibody, respectively, followed by BDNF addition does not prevent its effect ($N$ = 4 rats per condition). Results of statistical analyses are shown in Extended Data Fig. 5e. * $p < 0.05$ vs. vehicle (Veh), b $p < 0.05$ vs. ANA12 + anti-p75.

**a**

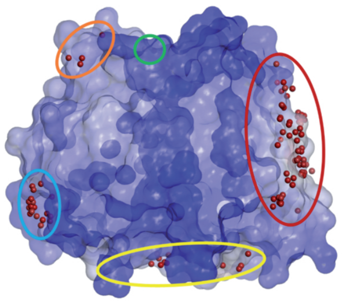

**c**

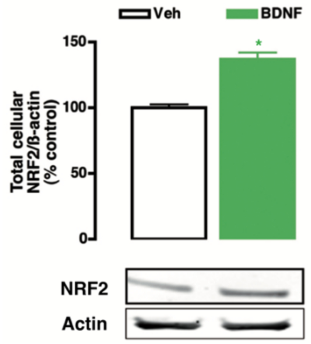


**b**

| Regions (ordered by no. poses) | Main clusters (Top 10) | Surrounding clusters | Number of poses (total poses 1620) | Poses in top 200 |
|---|---|---|---|---|
| Region 1 | 3 / 5 | 11 / 17 / 19 / 21 / 23 / 24 / 26 / 33 / 34 / 40 / 42 / 43 / 48 / 61 / 65 / 67 / 74 / 76 / 77 / 79 / 88 / 89 / 100 | 433 (26.7%) | 33 (16.5%) |
| Region 2 | 2 / 4 / 8 / 10 | 18 / 31 / 37 / 38 / 55 / 72 / 81 / 82 / 86 / 87 / 90 / 92 / 96 | 306 (18.9%) | 56 (28%) |
| Region 3 | 1 | 35 / 44 / 60 | 107 (6.6%) | 0 (0%) |
| Region 4 | 6 | 14 / 16 / 97 / 99 | 101 (6.3%) | 23 (11.5%) |
| Region 5 | 7 | 15 / 30 / 85 / 91 / 95 | 101 (6.3%) | 18 (9%) |

**Extended data Fig. 2 | BDNF interacts with KEAP1. a,** Side view of the KEAP1 solvent-accessible surface area (with transparence and colored according to pose concentration on its surface) and the top 100 poses of the 10 largest clusters. Yellow circle for Region 1, red circle for Region 2, orange circle for Region 3, blue circle for Region 4, and green circle for Region 5. **b,** Region analysis for the interaction between KEAP1 and BDNF. The main cluster centers are specified, along with the surrounding clusters populating the region. The percentage of total poses was calculated for classification, and the percentage of the top 200 poses for assessing the quality of the docking regions. **c,** Immunoblotting for NRF2 protein levels in cell lysate of HT-22 cells treated with BDNF or vehicle (Veh). Total cellular NRF2 protein levels increase in lysates following BDNF treatment (*N* = 3 experiments). * $p < 0.05$ vs. vehicle (Veh).

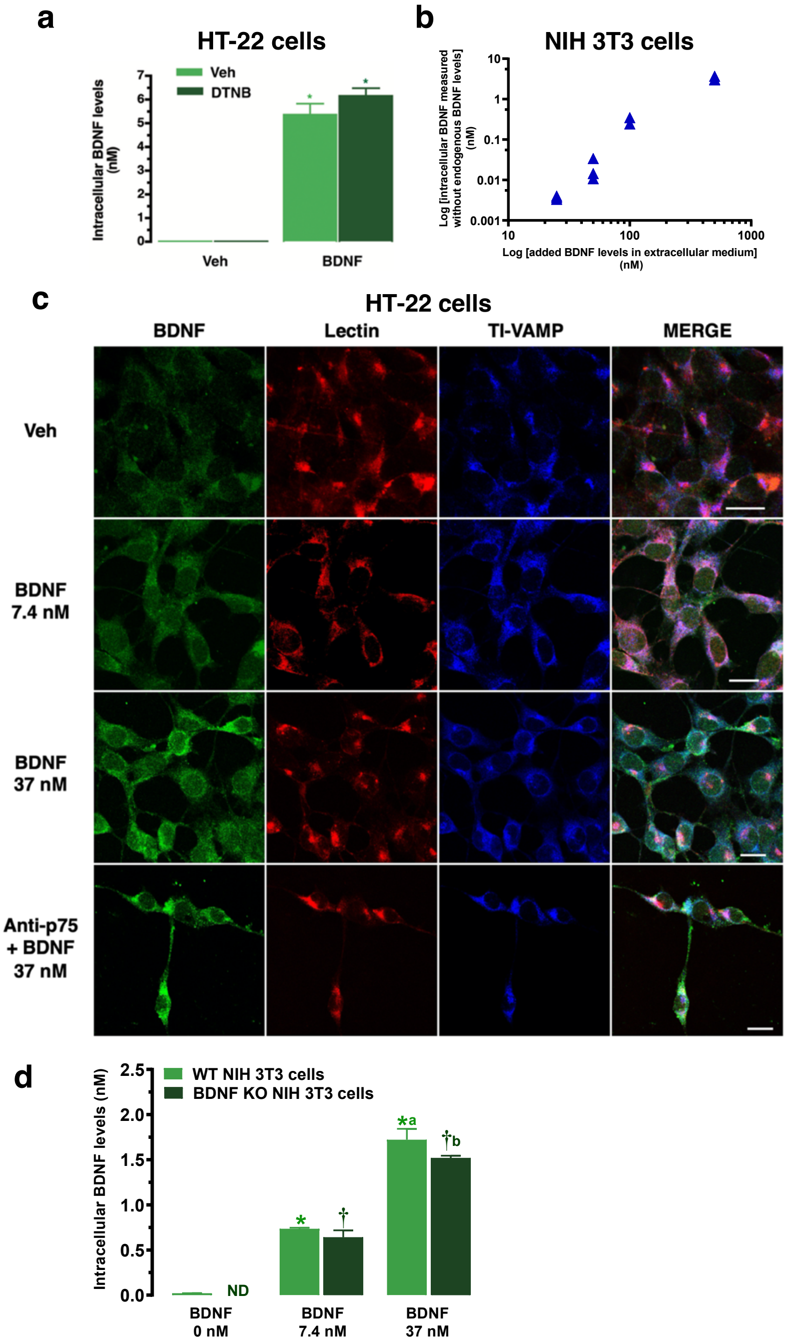


**Extended Data Fig. 3 | See next page for caption.**

**Extended Data Fig. 3 | BDNF crosses the cell membrane. a,** BDNF quantification on the cytosolic fraction of HT-22 cells treated with vehicle (Veh) or BDNF (37 nM). Addition of DTNB (2.5 mM), an inhibitor of protein-disulfide isomerase (PDI), followed by addition of BDNF does not modify the intracellular increase in BDNF induced by extracellular BDNF supplementation. Intracellular BDNF levels are expressed per 6.0 x $10^6$ - 6.5 x $10^6$ cells ($N = 2$ experiments per condition). Results of statistical analyses are shown in Extended Data Fig. 5e. * $p < 0.05$ vs. vehicle (Veh). **b,** BDNF quantification on the intracellular fraction of NIH 3T3 cells treated with BDNF (1.85-74 nM). Incubation with increasing concentrations of BDNF (1.85-37 nM) induce a linear increase in intracellular BDNF levels in NIH 3T3 cells, which do not express any BDNF known receptors. Intracellular BDNF levels are expressed per 5.0 x $10^6$ - 5.5 x $10^6$ cells ($N = 3$ experiments). **c,** Immunofluorescence experiments with BDNF antibody (green), lectin (WGA, red) and TI-VAMP antibody (blue) on HT-22 cells treated with BDNF (7.4 or 37 nM). Immunofluorescence experiments show a concentration-dependent increase of intracellular BDNF. Blockade of $p75^{NTR}$ receptors with a $p75^{NTR}$ -blocking antibody, followed by addition of BDNF (37 nM) does not affect its intracellular accumulation. Scale bar: 20 µm. **d,** BDNF quantification on the intracellular fraction of WT NIH 3T3 and BDNF KO NIH 3T3 cells treated with BDNF (0, 7.4 or 37 nM) at 4°C. Incubation with increasing concentrations of BDNF induce an increase in intracellular BDNF levels in Wild Type (WT) NIH 3T3 and in BDNF KO NIH 3T3 cells, which do not express BDNF in basal conditions (ND: non detectable). Intracellular BDNF levels are expressed per 5.0 x $10^6$ - 5.5 x $10^6$ cells ($N = 4$ experiments). Results of statistical analyses are shown in Extended Data Fig. 5e. * $p < 0.05$ vs. WT BDNF 0 nM, † $p < 0.05$ vs. BDNF KO BDNF 0 nM, a $p < 0.05$ vs. WT BDNF 7.4 nM, b $p < 0.05$ vs. BDNF KO BDNF 7.4 nM.

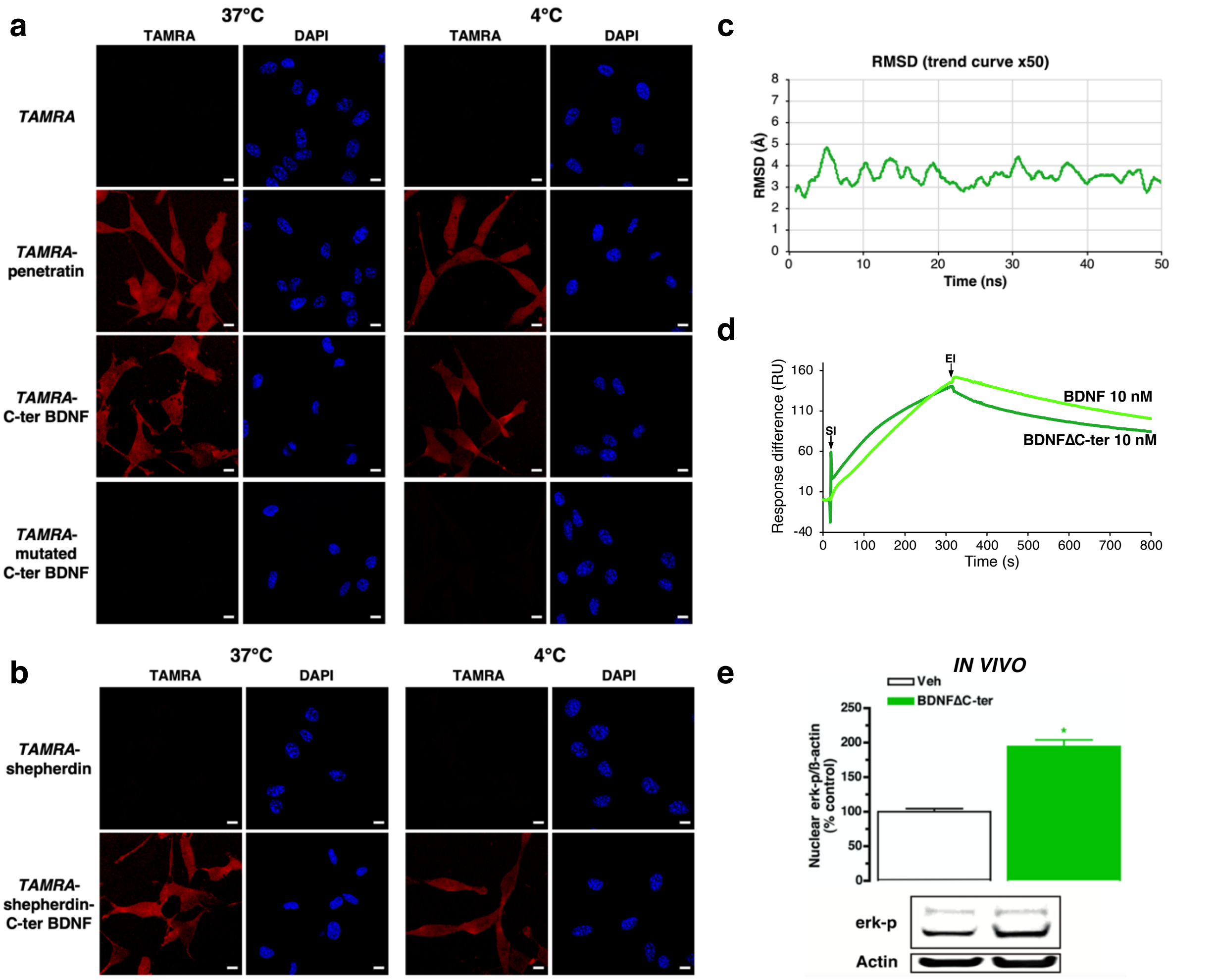


**Extended Data Fig. 4 | C-ter BDNF is necessary and sufficient to cross the cell membrane. a,** Confocal microscopy of NIH 3T3 cells incubated with *TAMRA*-labeled C-ter BDNF (1 µM), with *TAMRA*-labeled penetratin as positive control (1 µM), with *TAMRA* fluorophore (1 µM), an impermeant cargo, or with *TAMRA*-mutated C-ter BDNF (1 µM) during 20 minutes at 37°C and at 4°C. Cell nuclei were stained with DAPI. Extracellular application of *TAMRA*-labeled C-ter BDNF to NIH 3T3 cells resulted in its accumulation in the cytosol, as *TAMRA*-labeled penetratin. *TAMRA* fluorophore does not accumulate intracellularly. *TAMRA*-mutated C-ter BDNF does not accumulate intracellularly ($N$ = 6 experiments per condition). Scale bar: 10 µm. **b,** Confocal microscopy of NIH 3T3 cells incubated with *TAMRA*-shepherdin, an impermeant peptide, conjugated or not to C-ter BDNF (1 µM). Extracellular application of *TAMRA*-shepherdin-C-ter BDNF to NIH 3T3 cells results in its accumulation in the cytosol. *TAMRA*-shepherdin does not lead to any intracellular accumulation. Each experiment has been performed at 37°C and at 4°C ($N$ = 6 experiments per condition). Scale bar: 10 µm. **c,** Root mean square deviation of BDNFΔC-ter against the first frame of the simulation. Interestingly, the structure of BDNFΔC-ter underwent minor adjustments (RMSD of 3 Å) at the beginning of the simulation, then remained very stable throughout the 50 ns. **d,** Surface plasmon resonance (SPR) was performed between KEAP1 and BDNF (10 nM) or BDNFΔC-ter (10 nM). SI: start of injection; EI: end of injection. Sensorgrams demonstrate a direct interaction between KEAP1 and BDNFΔC-ter with a similar signal and identical curve profile as BDNF ($N$ = 3 experiments). **e,** Immunoblotting for phospho ERK (erk-p) on a nuclear extract from rat hippocampus after *in vivo* infusion of BDNFΔC-ter (0.8 µg) or vehicle (Veh). Infusion of BDNFΔC-ter into the rat hippocampus activates the ERK pathway ($N$ = 3 rats per condition). * $p < 0.05$ vs. vehicle (Veh).

**a**

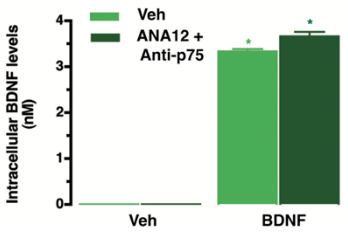


**b**

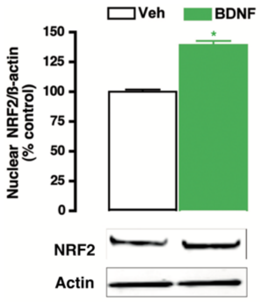


**c**

| |
|---|
| Amphiregulin |
| Bone morphogenetic protein 15 |
| Epidermal growth factor receptor kinase substrate 8-like protein 2 |
| Fibroblast growth factors: FGF2, FGF3, FGF12, FGF13, FGF14 |
| Fibroblast growth factor-binding protein 3 |
| Growth arrest-specific protein 6 |
| Growth/differentiation factors : GDF5, GDF6, GDF10 |
| Latent-transforming growth factor beta-binding protein 4 |
| Placenta growth factor |
| Pleiotrophin |
| Vascular endothelial growth factors : VEGF A , VEGF D |

**d**

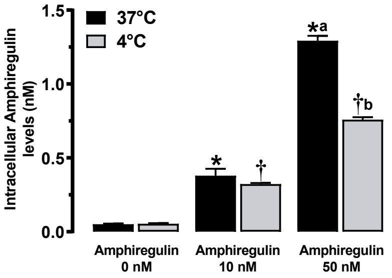


**e**

| **Fig. 1** | Panel B : Nuclear NRF2 HT-22 cells | Panel B : Cytosolic NRF2 HT-22 cells | Panel E : Nuclear NRF2 NIH 3T3 cells |
|---|---|---|---|
| Treatment effect | $F(2, 9) = 116.6$; $p < 0.0001$ | $F(2, 9) = 48.17$; $p < 0.0001$ | $F(2, 6) = 16.81$; $p = 0.003$ |

| **Fig. 3** | Panel A : Intracellular BDNF WT/BDNF KO NIH 3T3 cells |
|---|---|
| Group effect | $F(1, 6) = 1.392$; $p = 0.28$ |
| Treatment effect | $F(1.056, 6.339) = 164.3$; $p < 0.0001$ |

| **Fig. 4** | Panel D : Nuclear NRF2 HT-22 cells |
|---|---|
| Treatment effect | $F(2, 9) = 38.01$; $p < 0.0001$ |

| **Extended Data Fig. 1** | Panel C : Cytosolic KEAP1 HT-22 cells | Panel D : Nuclear p-ERK HT-22 cells | Panel E : Nuclear NRF2 in vivo |
|---|---|---|---|
| Treatment effect | $F(2, 9) = 2.56$; $p = 0.13$ | $F(2, 6) = 36.58$; $p = 0.0004$ | $F(3, 12) = 15.5$; $p = 0.0002$ |

| **Extended Data Fig. 3** | Panel A : Intracellular BDNF HT-22 cells | Panel D : Intracellular BDNF WT/BDNF KO NIH 3T3 cells |
|---|---|---|
| Group effect | $F(1, 26) = 569.90$; $p < 0.0001$ | $F(1, 6) = 5.439$; $p = 0.058$ |
| Treatment effect | $F(1, 26) = 2.79$; $p = 0.10$ | $F(2, 12) = 345.2$; $p < 0.0001$ |
| Group x Treatment | $F(1, 26) = 2.78$; $p = 0.10$ | |

| **Extended Data Fig. 5** | Panel A : Intracellular BDNF Caco-2 cells | Panel D : Intracellular amphiregulin HT-22 cells at 37°C or 4°C |
|---|---|---|
| Group effect | $F(1, 8) = 4078$; $p < 0.0001$ | $F(1, 6) = 220.7$; $p < 0.0001$ |
| Treatment effect | $F(1, 8) = 9.01$; $p = 0.01$ | $F(1.273, 7.638) = 636.7$; $p < 0.0001$ |
| Group x Treatment | $F(1, 8) = 8.99$; $p = 0.01$ | $F(2, 12) = 55.42$; $p < 0.0001$ |

**Extended Data Fig. 5 | See next page for caption.**

**Extended Data Fig. 5 | BDNF crosses the cell membrane in differentiated intestinal epithelial cells – CPP-like sequences in other proteins. a,** BDNF quantification by an ELISA Assay on the cytosolic fraction of Caco-2 intestinal cells treated with BDNF (7.4 nM) or vehicle (Veh). Extracellular addition of BDNF induces its intracellular accumulation in Caco-2 cells. Blockade of TrkB and $p75^{NTR}$ receptors with the ANA12 inhibitor and a $p75^{NTR}$-blocking antibody, respectively, followed by BDNF addition did not prevent intracellular accumulation. Intracellular BDNF levels are expressed per 2.5 x $10^5$ cells ($N$ = 3 experiments). Results of statistical analyses are shown in Extended Data Fig. 5e. * $p < 0.05$ vs. vehicle (Veh). **b,** Immunoblotting for NRF2 protein levels in nuclear fraction of Caco-2 cells treated with BDNF (7.4 nM) or vehicle (Veh). Incubation of Caco-2 cells with BDNF induces an increase in nuclear NRF2. Blockade of TrkB and $p75^{NTR}$ receptors with the ANA12 inhibitor and a $p75^{NTR}$-blocking antibody, respectively, followed by BDNF addition does not prevent NRF2 nuclear translocation ($N$ = 3 experiments). * $p < 0.05$ vs. vehicle (Veh). **c,** *In silico* analysis of CPP-like sequences in growth factors and associated proteins. Eighteen proteins, mainly growth factors, possess a CPP-like sequence, which may allow them to cross the cell membrane. Whether or not they do the latter, as well as their intracellular targets, remain to be determined. **d,** Amphiregulin quantification on the intracellular fraction of HT-22 cells treated with amphiregulin (0, 10, 50 nM) at 37°C and 4°C. Incubation with extracellular amphiregulin induces an increase in intracellular amphiregulin levels at 37°C and 4°C. Intracellular amphiregulin levels are expressed per 6.0 x $10^6$ - 6.5 x $10^6$ cells ($N$ = 4 experiments). Results of statistical analyses are shown in Extended Data Fig. 5e. * $p < 0.05$ vs. amphiregulin 0 nM 37°C, † $p < 0.05$ vs. amphiregulin 0 nM 4°C, a $p < 0.05$ vs. amphiregulin 10 nM 37°C, b $p < 0.05$ vs. amphiregulin 10 nM 4°C. **e,** Table showing results of statistical analyses**. Fig. 1:** Panel b: Statistical analysis of the effects of either BDNF or quercetin on nuclear and cytosolic NRF2 concentrations in HT-22 cells, based on one-way ANOVA. Panel e: Statistical analysis of BDNF effects (0, 7.4 or 37 nM) on nuclear NRF2 concentrations in NIH 3T3 cells, based on one-way ANOVA. **Fig. 3:** Panel a: Statistical analysis of the effect of BDNF or vehicle on intracellular BDNF levels, in WT or BDNF KO NIH 3T3 cells at 37°C, based on two-way ANOVA. **Fig. 4:** Panel d: Statistical analysis of the effects of either BDNF or BDNF∆CPP on nuclear NRF2 concentrations in HT-22 cells, based on one-way ANOVA. **Extended Data Fig. 1:** Panel c: Statistical analysis of the effects of either BDNF or quercetin on cytosolic Keap1 concentrations in HT-22 cells, based on one-way ANOVA. Panel d: Statistical analysis of BDNF effect on p-ERK concentration, in HT-22 cells, based on one-way ANOVA. Panel e: Statistical analysis of BDNF effect of BDNF on nuclear NRF2 concentration, in *in vivo* conditions, based on one-way ANOVA. **Extended Data Fig. 3:** Panel a: Statistical analysis of the effects of either DTNB or vehicle and of either BDNF or vehicle on intracellular BDNF levels, in HT-22 cells, based on two-way ANOVA. Panel d: Statistical analysis of the effect of either BDNF or vehicle on intracellular BDNF levels, in either WT or BDNF KO NIH 3T3 cells at 4°C, based on two-way ANOVA. **Extended data Fig. 5:** Panel a: Statistical analysis of the effects of either ANA12 + Anti-p75 or vehicle and either BDNF or vehicle on intracellular BDNF levels, in Caco-2 cells, based on two-way ANOVA. Panel d: Statistical analysis of the effect of either amphiregulin or vehicle on intracellular amphiregulin levels, in HT-22 cells at 37°C or 4°C, based on two-way ANOVA.

### Supplementary Information

**Supplementary Video 1 | Interaction between BDNF and KEAP1.** The first scene features a side view of KEAP1. The second features a side view of BDNF. The third and fourth scenes feature the best poses of BDNF in Regions 1 and 2, identified as the best hypothetical binding regions. The solvent-accessible surface area is colored according to its hydrophobicity, from blue (hydrophilic) to brown (hydrophobic). The secondary structure of BDNF is colored red for the alpha helix, and blue for the beta sheet. The dots represent the center of mass for the interface residues of the poses, colored from blue (lowest score) to red (highest score).

**Supplementary Video 2 | Internalization of *TAMRA*-C-ter BDNF in HT-22 cells.** Live-cell imaging was performed on HT-22 cells incubated with *TAMRA*-C-ter BDNF (*TAMRA*-KKRIGWRFIRIDTSCVCTLTIKRGR-COOH; 1 µM) in an incubation chamber at 37 °C in a 5% $CO_2$ atmosphere. Acquisition was performed every 30 seconds for 40 minutes ($N$ = 6 experiments). After washing, Differential Interference Contrast (DIC) imaging was performed.
